# Gaze patterns and brain activations in humans and marmosets in the Frith-Happé theory-of-mind animation task

**DOI:** 10.1101/2023.01.16.524238

**Authors:** Audrey Dureux, Alessandro Zanini, Janahan Selvanayagam, Ravi S. Menon, Stefan Everling

**Affiliations:** Centre for Functional and Metabolic Mapping, Robarts Research Institute, University of Western Ontario, London, ON N6A 5K8, Canada; Department of Physiology and Pharmacology, University of Western Ontario, London, ON N6A 5K8, Canada

**Author notes:** <u>Corresponding author:</u> Audrey Dureux, Centre for Functional and Metabolic Mapping, Robarts Research Institute, University of Western Ontario, London, Canada.

**Keywords:** Theory of mind, Frith-Happé animations, fMRI, eye tracking, awake marmosets

## Abstract

Theory of Mind (ToM) refers to the ability to ascribe mental states to other individuals. This process is so strong that it extends even to the attribution of mental states to animations depicting interacting simple geometric shapes, such as in the Frith-Happé animations in which two triangles move either purposelessly (Random condition), or as if one triangle is reacting to the other triangle’s mental state (ToM condition). Currently, there is no evidence that nonhuman primates attribute mental states to moving abstract shapes. Here we investigated whether highly social marmosets (*Callithrix jacchus*) process ToM and Random Frith-Happé animations differently. Our results show that marmosets and humans (1) follow more closely one of the triangles during the observation of ToM compared to Random animations, and (2) activate large and comparable brain networks when viewing ToM compared to Random animations. These findings indicate that marmosets, like humans, process ToM animations differently from Random animations.

## Introduction

Theory of Mind (ToM) refers to the capacity to ascribe mental states to other subjects (Carruthers and Smith, 1996; Premack and Woodruff, 1978). A variety of experimental approaches have been developed to study the cognitive processes underlying ToM, such as text-based tasks (Happé, 1994), non-verbal picture-based tasks (Sarfati et al., 1997), false belief tasks (Wimmer and Perner, 1983), and silent animations of geometric shapes. The latter approach is based on Heider and Simmel’s observation that participants attribute intentional actions, human character traits, and even mental states to moving abstract shapes (Heider and Simmel, 1944). Subsequent studies used these animations to test the ability to ascribe mental states in autistic children (Bowler and Thommen, 2000; Klin, 2000).

In the popular computer-generated Frith-Happé animations, a large red triangle and a small blue triangle move around the screen (Abell et al., 2000; Castelli et al., 2002, 2000). In the Random condition, the two triangles do not interact and move about purposelessly, whereas in the ToM condition the two animated triangles move as if one triangle is reacting to the other’s mental state (i.e., coaxing, surprising, seducing and mocking). Functional imaging studies have demonstrated that the observation of ToM compared to Random animations activates brain regions typically associated with social cognition, including medial frontal, temporoparietal, inferior and superior temporal cortical regions (Barch et al., 2013; Castelli et al., 2000; Wheatley et al., 2007).

While it has been now well established that humans spontaneously ascribe mental states to moving shapes, it is largely unknown whether other primate species also possess this capacity. There is some evidence that monkeys can attribute mental states to some moving stimuli (e.g., moving dots with apparent biological motion), but findings are mixed (Atsumi et al., 2017; Atsumi and Nagasaka, 2015; Krupenye and Hare, 2018; Kupferberg et al., 2013; Uller, 2004). The degree to which nonhuman primates spontaneously attribute mental states to inanimate objects is even less certain. Whereas human subjects have longer eye fixations when viewing the two triangles in the ToM condition than in the Random condition of the Frith-Happé animations, a recent eye tracking study did not find any differences in macaque monkeys (Schafroth et al., 2021).

Here, we investigated the behaviour and brain activations of New World common marmoset monkeys (*Callithrix jacchus)* while they watched Frith-Happé animations. In contrast to macaques, marmosets live in family groups and share important social similarities with humans, such as prosocial behavior, imitation, and cooperative breeding, making them a promising nonhuman primate model for the study of social cognition (Burkart et al., 2009; Burkart and Finkenwirth, 2015; Miller et al., 2016). To directly compare humans and marmosets while viewing these animations, we used high-speed video eye tracking to measure saccades and fixations in twelve healthy humans and twelve marmoset monkeys and we acquired ultra-high field fMRI data in ten healthy humans at 7T and six common marmoset monkeys at 9.4T. The results indicate that marmosets, similar to humans, process ToM and Random animations differently, and activate similar brain networks when viewing ToM compared to Random animations.

## Results

To investigate whether marmoset monkeys, like humans, process ToM and Random animations differently, we compared gaze patterns and fMRI activations while marmosets and human subjects watched shortened version of the Frith-Happé animations (Figure 1).

**Figure 1.**
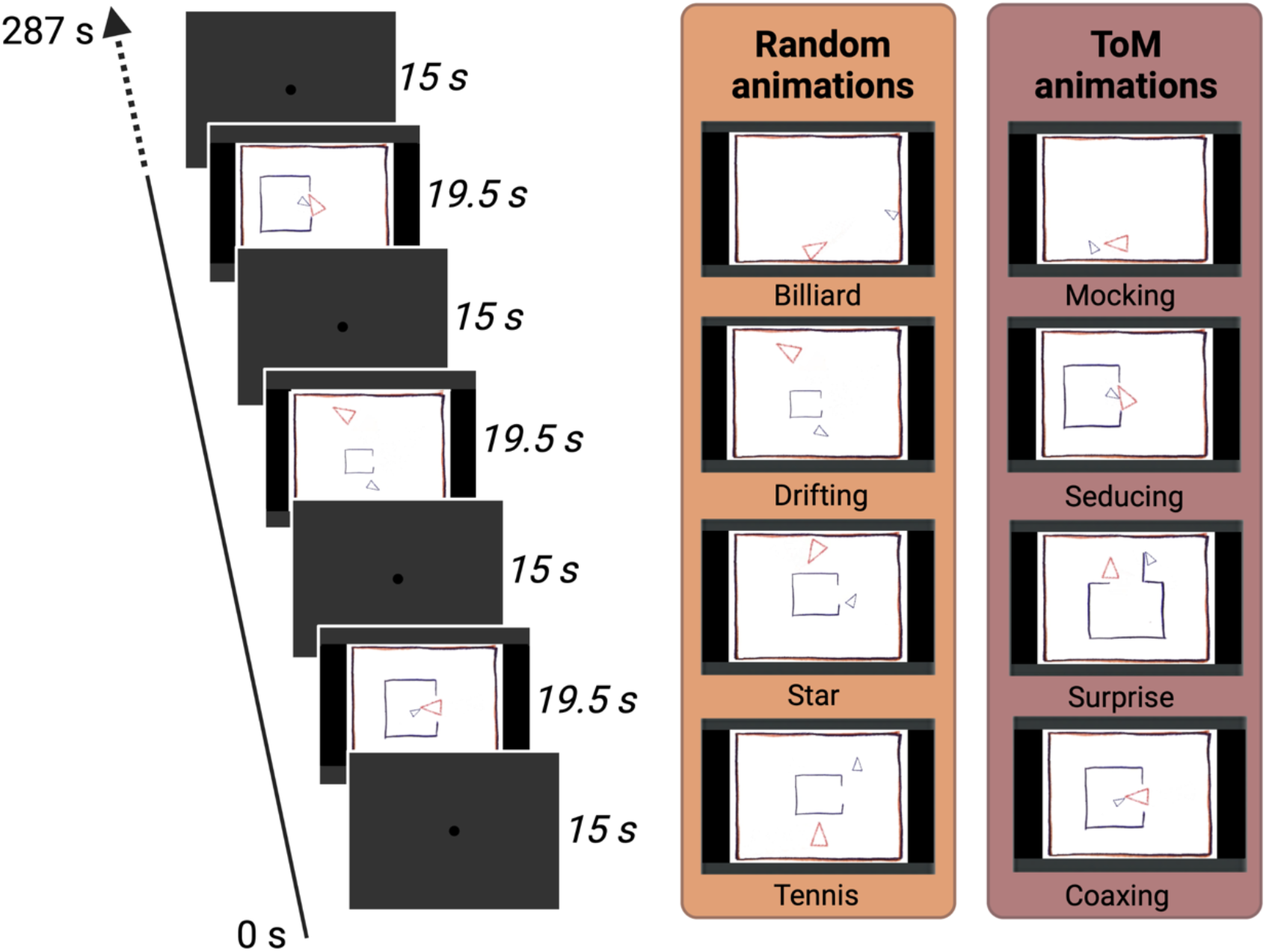
Task Design. In each run, two different types of video clips were presented four times each in a Randomized order. In the ToM animations, one triangle reacted to the other triangle’s mental state, whereas in the Random animations the same two triangles didn’t interact with each other. Each animation video lasted 19.5 sec and was separated by baseline blocks of 15 sec where a central dot was displayed in the center of the screen. In the fMRI task, several runs were used with a Randomized order of the conditions whereas in the eye-tracking task one run containing all the eight conditions once was used.

### Gaze patterns for Frith-Happé’s ToM and Random animations in humans and marmosets

We first investigated in both humans and marmosets whether fixation durations differed between ToM and Random conditions. By conducting mixed analyses of variance (ANOVA), with factors of species (Human vs Marmoset) and condition (ToM vs Random animation videos), we found a significant interaction between species and condition (*F*_(1,22)_=7.67, *p*=.01, *η*_*p*_^*2*^=.258). Here we observed longer fixation durations for ToM animation videos (*M*=317.2 ms) as compared to Random videos (*M*=269.0 ms) for humans (*p* =.029) but not for marmosets (258.3 ms vs 270.5 ms, *p*=.81).This finding confirms that humans fixate longer in the ToM condition (Klein et al., 2009) whereas marmosets, like macaques (Schafroth et al., 2021), do not show this effect.

To examine the gaze patterns of humans and marmosets in more detail, we next measured the proportion of time subjects looked at each of the triangles in the videos (Figure 2). We conducted mixed ANOVAs on the proportion of time the radial distance between the current gaze position and each triangle was within 4 visual degrees for each triangle separately. For the large red triangle, we observed a significant interaction of species and condition (*F*_(1,22)_=21.4, *p*<.001, *η*_*p*_^*2*^ =.493). Both humans (Figure 2 left; Δ=.317, *p*<.001) and marmosets (Figure 2 right; Δ=.145, *p*<.001) spent a greater proportion of time looking at the red triangle in TOM compared to Random videos. For the small blue triangle, we also observed a significant interaction of species and condition (*F*_(1,22)_=10.7, *p*=.003, *ηp2*=.328) but only humans (Δ=.125, *p*=.001). They spent a significantly greater proportion of time looking at the blue triangle in ToM than in Random animation videos, whereas no significant differences were observed for marmosets (Δ=.002, *p*>.999; Figure 2). These results demonstrate that the eye tracking patterns of both humans and marmosets varies between ToM and Random videos.

**Figure 2.**
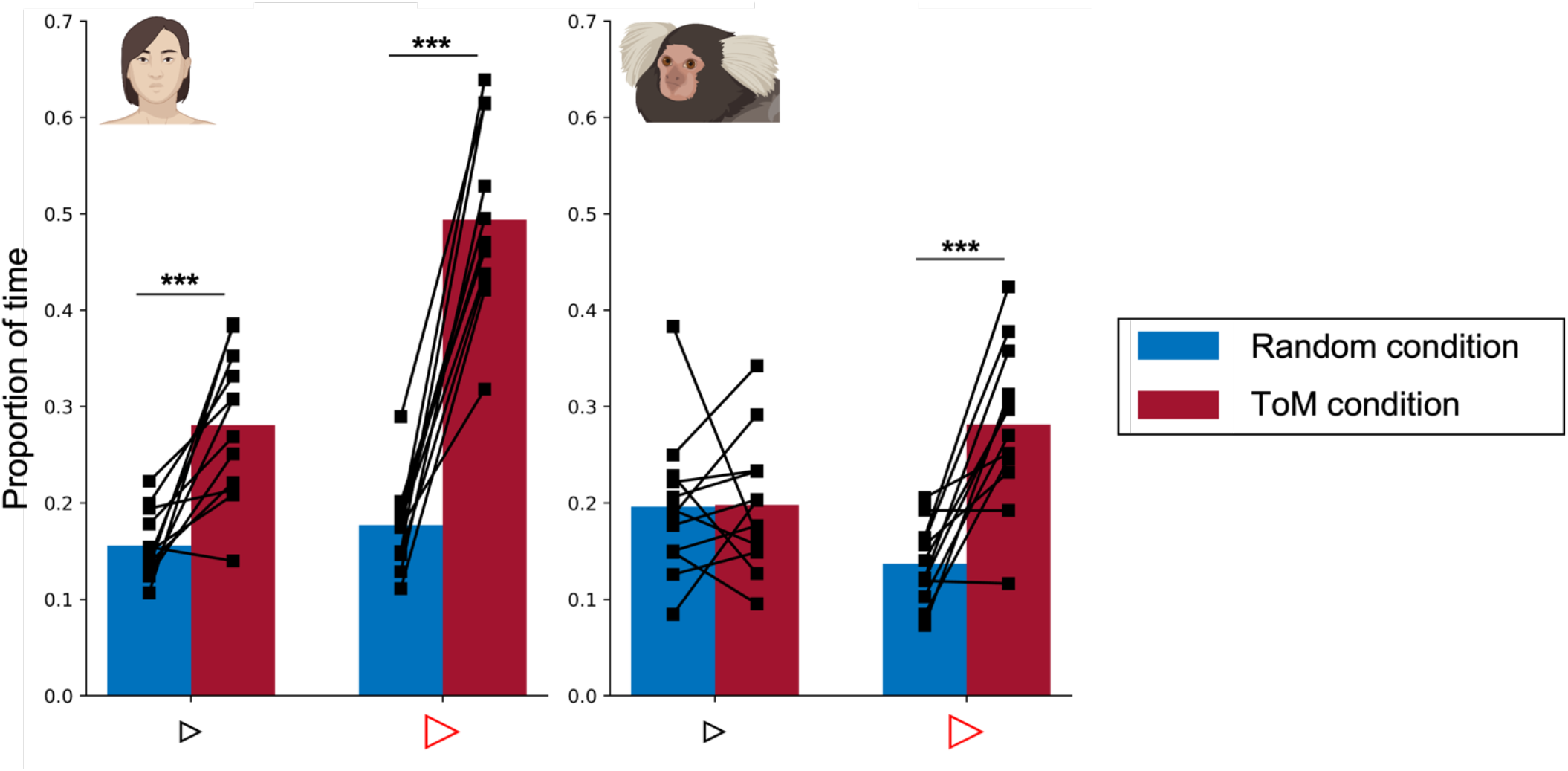
Proportion of time looking triangles in Frith-Happé’s ToM and Random animations in humans (left) and marmosets (right). Bar plot representing the proportion of time the radial distance between the current gaze position and each triangle was within 4 visual degrees, as a function of each condition. Red represents results obtained for ToM animation videos and blue represents results for Random animation videos. The left panel shows the results for 12 humans and the right panel for 12 marmosets. Each colored bar represents the group mean and the thick black lines represent individual results. The differences between conditions were tested using ANOVA: *p*<.05*, *p*<.01** and *p*<.001***.

### Functional brain activations while watching ToM and Random Frith-Happé’s animations in humans

We first investigated ToM and Random animations processing in humans. Figure 3 shows group activation maps for ToM (A) and Random (B) conditions as well as the comparison between ToM and Random conditions (C) obtained for human participants.

**Figure 3.**
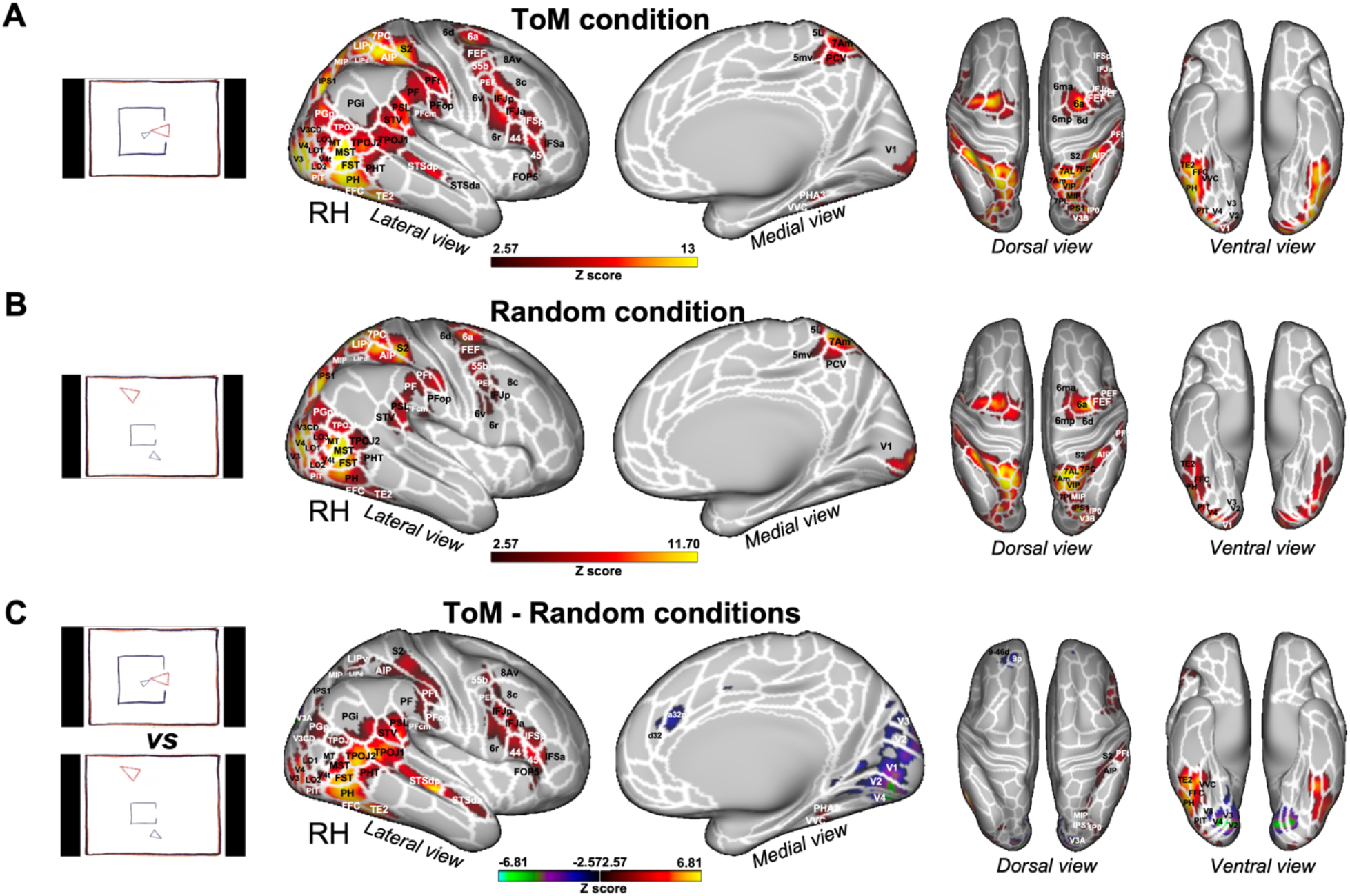
Brain networks involved in processing of Frith-Happé’s ToM and Random animations in humans. Group functional maps displayed on right fiducial (lateral and medial views) and left and right fiducial (dorsal and ventral views) of human cortical surfaces showing significant greater activations for ToM condition (A), Random condition (B) and the comparison between ToM and Random conditions (C). The white line delineates the regions based on the recent multi-modal cortical parcellation atlas (Glasser et al., 2016). The maps depicted are obtained from 10 human subjects with an activation threshold corresponding to z-scores > 2.57 for regions with yellow/red scale or z-scores < -2.57 for regions with purple/green scale (AFNI’s 3dttest++, cluster-forming threshold of p<.01 uncorrected and then FWE-corrected α=.05 at cluster-level from 10000 Monte-Carlo simulations).

Both ToM (Figure 3A) and Random (Figure 3B) videos activated a large bilateral network including visual areas (V1, V2, V3, V3CD, V3B, V4, V4T, V6A, V7, MT, MST), lateral occipital areas 1, 2 and 3 (LO1, LO2, LO3), temporal areas (FST, PH, PHT, TE2, posterior inferotemporal complex PIT and fusiform face complex FFC), temporo-parietal junction areas (TPOJ2 and TPOJ3), lateral posterior parietal areas also comprising the parietal operculum (supramarginal areas PF, PFt, PFop and PFcm, angular areas PGp and PGi, superior temporal visual area STV, perisylvian language area PSL, medial intraparietal area MIP, ventral and dorsal lateral intraparietal areas LIPv and LIPd, anterior intraparietal area AIP, IPS1, IPS0, 7PC and 5L), medial superior parietal areas (7am, PCV, 5mv), secondary somatosensory cortex (S2), premotor areas (6, 55b, premotor eye field PEF, frontal eye field FEF), and frontal areas (8C, IFJp).

The ToM condition (Figure 3A) also showed bilateral activations in posterior and anterior superior temporal sulcus (STSdp and STSa), in temporo-parietal junction area TPOJ1, in ventral visual complex (VVC), in parahippocampal area 3 (PHA3), in inferior part of angular area (PGi), in lateral prefrontal areas 8C and 8Av, in inferior frontal areas IFJa, IFSp, IFSa and in frontal opercular area 5 (FOP5).

To identify brain areas that are more active during the observation of ToM compared to Random videos, we directly compared the two conditions (i.e., ToM animations > Random animations contrast, Figure 3C and Figure 5A). This analysis shows stronger activations for the ToM condition in occipital and temporal areas with significant differences in bilateral visual areas V3, V3CD, V4, V4t, MT, MST, in bilateral LO1 and LO2, within the lateral temporal lobe in bilateral areas PHT, PH, FST and in the more inferior part of the temporal lobe in bilateral areas TE2, FFC, PIT, in bilateral temporo-parietal junction areas TPOJ1, TPOJ2, TPOJ3 and along the right STS in STSdp and STSda areas. We also observed stronger activations in left and right parietal areas, in the inferior parietal lobule (right supramarginal and opercular supramarginal areas PF, PFt, PFop and PFcm, bilateral opercular areas PSL and STV, bilateral angular areas PGp and PGi, right IPS1 and right MIP) and in the superior parietal lobule (right AIP, LIPv and LIPd). More anteriorly we found greater activations in the right hemisphere in secondary somatosensory cortex, premotor areas (55b, 6r and PEF, right hemisphere), lateral prefrontal areas (8C, 8Av, 44 and 45, right hemisphere), inferior (IFSa, IFSp, IFJa and IFJp, right hemisphere) and opercular (FOP2, FOP5) frontal areas. Stronger activations for the Random condition were mainly limited to left and right visual areas (V1, V2, V3, V3A, V4, V8), as well as in lateral (9p bilateral, 9-46d and 8Ad left) and medial frontal areas (a32p and 24dv bilateral, d32 right, and a32pr left).

**Figure 4.**
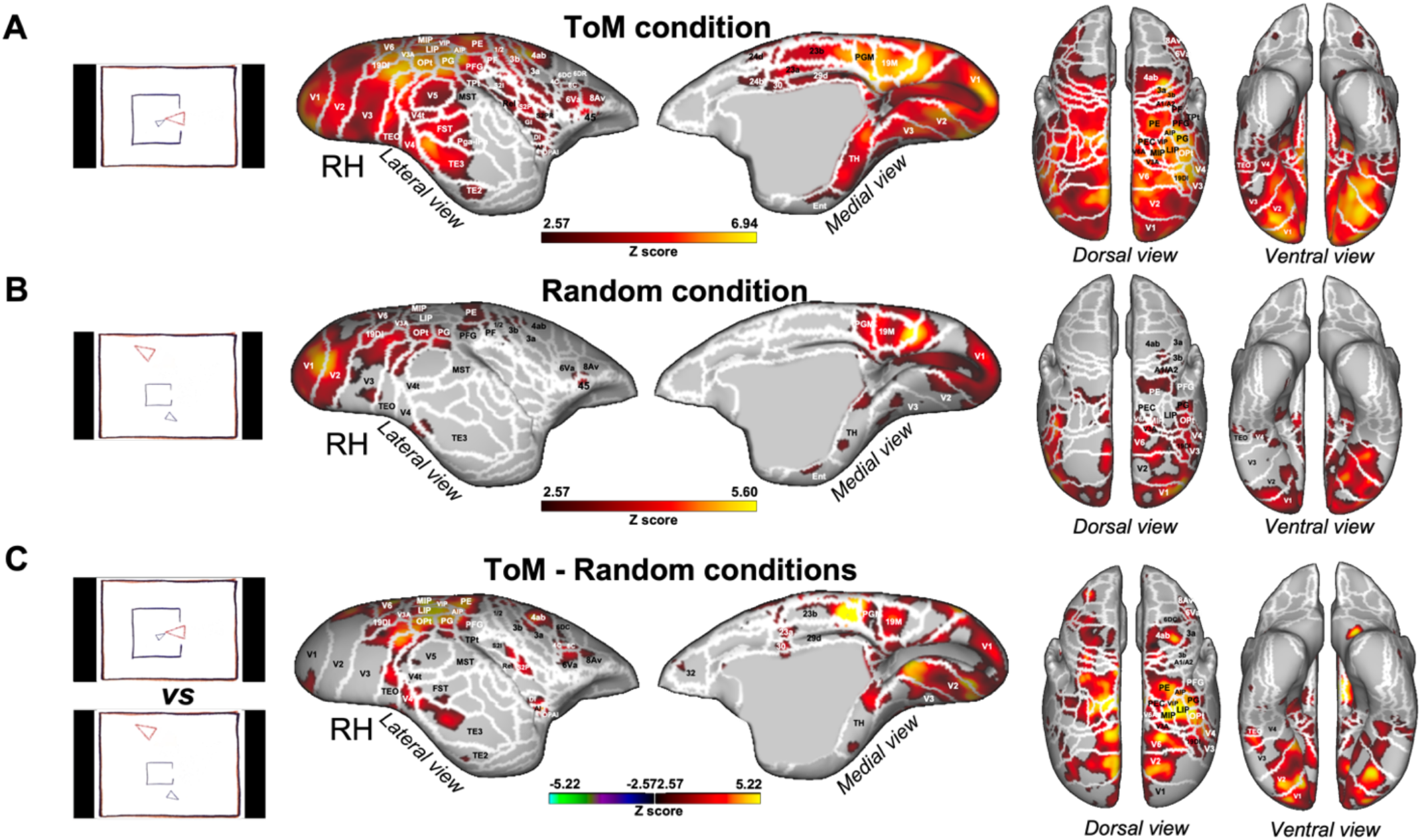
Brain networks involved in processing of Frith-Happé’s ToM and Random animations in marmosets. Group functional maps showing significant greater activations for ToM condition **(A)**, Random condition **(B)** and the comparison between ToM and Random conditions **(C)**. Group map obtained from 6 marmosets displayed on lateral and medial views of the right fiducial marmoset cortical surfaces as well as dorsal and ventral views of left and right fiducial marmoset cortical surfaces. The white line delineates the regions based on the Paxinos parcellation of the NIH marmoset brain atlas (Liu et al., 2018). The brain areas reported have activation threshold corresponding to z-scores > 2.57 (yellow/red scale) or z-scores < -2.57 (purple/green scale) (AFNI’s 3dttest++, cluster-forming threshold of p<.01 uncorrected and then FWE-corrected α=.05 at cluster-level from 10000 Monte-Carlo simulations).

**Figure 5.**
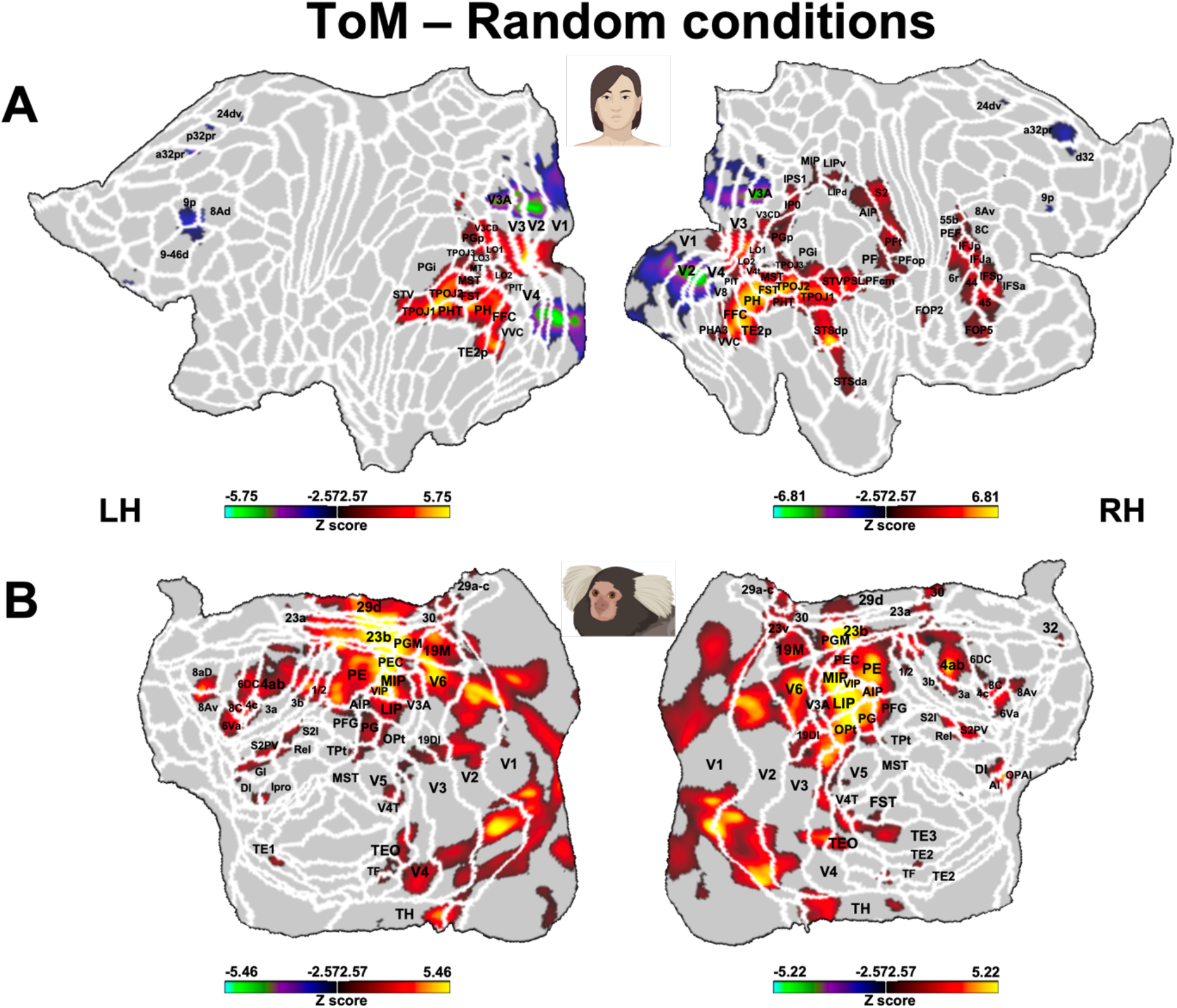
Brain network involved during processing of ToM compared to Random Frith-Happé’s animations in both humans (A) and marmosets (B). Group functional maps showing significant greater activations for ToM animations compared to Random animations. **A**. Group map obtained from 10 human subjects displayed on the left and right human cortical flat maps. The white line delineates the regions based on the recent multi-modal cortical parcellation atlas (Glasser et al., 2016). **B**. Group map obtained from 6 marmosets displayed on the left and right marmoset cortical flat maps. The white line delineates the regions based on the Paxinos parcellation of the NIH marmoset brain atlas (Liu et al., 2018). The brain areas reported in A and B have activation threshold corresponding to z-scores > 2.57 (yellow/red scale) or z-scores < -2.57 (purple/green scale) (AFNI’s 3dttest++, cluster-forming threshold of p<.01 uncorrected and then FWE-corrected α=.05 at cluster-level from 10000 Monte-Carlo simulations).

At the subcortical level (see Supplementary Figure S1 left panel), we found stronger bilateral activations in the cerebellum, in some portions of the thalamus (in right ventroposterior thalamus THA-VP and left and right dorsoanterior thalamus THA-DA) and amygdala for both ToM (Supplementary Figure S1A left panel) and Random conditions (Supplementary Figure S1B left panel). Activations were stronger in posterior lobe of cerebellum, right amygdala and thalamus (right THA-VP, right ventroanterior thalamus THA-VA and left and right dorsoposterior thalamus THA-DP) for the ToM condition compared to the Random condition and in the left and right side of the cerebellar cortex for Random compared to ToM conditions (Supplementary Figure S1C left panel).

As we used shorter modified versions of the Frith-Happé animations (i.e., videos of 19.5 sec instead of 40 sec), we also validated our stimuli and our fMRI protocol by comparing the brain responses elicited by ToM animation videos - compared to Random animation videos - obtained in our group of 10 human subjects and those reported by the large group of humans (496) used in the social cognition task of the HCP (Barch et al., 2013), which also used shortened versions of the Frith-Happé animations.

This comparison is shown in Supplementary Figure S2. Overall, we observed similar distinct patterns of brain activations (Supplementary Figure S2A and S2B), including a set of areas in occipital, temporal, parietal and frontal cortices, as described previously (Figure 3C). The main differences were stronger activations in the left hemisphere in the HCP dataset. Therefore, these results show that our stimuli and our protocol are appropriate to investigate mental state attribution to animated moving shapes.

### Functional brain activations while watching ToM and Random Frith-Happé’s animations in marmosets

Having identified the dedicated brain networks for ToM and Random video processing in human subjects and validating our protocol, we then used these same stimuli in marmosets. Figure 4 shows the brain network obtained by the ToM condition (A), Random condition (B) and ToM compared to Random condition (C) in six marmosets.

ToM (Figure 4A) and Random (Figure 4B) animations activated an extended network comprising a set of areas in occipito-temporal, parietal and frontal areas. We found higher bilateral activations in occipital and temporal cortex, in visual areas V1, V2, V3, V3A, V4, V4t, V5, V6, MST, 19 medial part (19M) and dorsointermediate part (19DI), in ventral temporal area TH and enthorinal cortex, in lateral and inferior temporal areas TE3 and TEO. We also observed greater activations in posterior parietal cortex, in bilateral regions surrounding the intraparietal sulcus (IPS), in LIP, MIP, PE, PG, PFG, PF, V6A, PEC, in occipito-parietal transitional area (OPt) and in medial part of the parietal cortex (area PGM). More anterior, bilateral activations were present in areas 1/2, 3a, 3b of the somatosensory cortex, in primary motor area 4 parts a, b and c (area 4ab and 4c), in area 6 ventral part (6Va) of the premotor cortex and in frontal areas 45 and 8Av.

The ToM condition (Figure 4A) recruited a larger network, with also greater bilateral activations in areas V5, TE2, FST, Pga-IPa, temporoparietal transitional area (TPt), around the IPS in AIP and VIP, in internal part (S2I), parietal rostral part (S2PR) and ventral part (S2PV) of the secondary somatosensory cortex, in agranular insular cortex (AI), granular and dysgranular insular areas (GI and DI), retroinsular area (ReI) and orbital periallocortex (OPAI), as well as in premotor cortex in area 8 caudal part (8C), in area 6 dorsocaudal and dorsorostral parts (6DC, 6DR). Moreover, we also observed higher activations in posterior cingulate areas 23a, 23b, 29d, 30, 24d and 24b.

Next, we identified areas that showed different activations for ToM compared to Random animations (i.e., ToM condition > Random condition contrast, Figure 4C and Figure 5B). We found stronger bilateral activations for ToM condition in occipital areas V1, V2, V3, V3A, V4, V4t, V5, V6, 19DI, 19M, in temporal areas TH, TE2, TE3, FST, MST, TPt, in parietal areas LIP, MIP, VIP, AIP, PE, PG, PFG, OPt, V6A, PEC, in somatosensory cortex areas 1/2, 3a, 3b, S2I, S2PV, in parts of primary motor cortex areas 4ab and 4c, in lateral frontal areas 6DC, 8C, 6Va, 8Av, 8Ad left, in insular areas ReI, S2I, S2PV, DI, AI, in OPAI area, in medial frontal area 32 and posterior cingulate areas 23a, 23b, 29d, 30. We found no regions with stronger activations for Random compared to ToM animations.

At the subcortical level (see Supplementary Figure S1A right panel), greater activations for the ToM condition were present in the bilateral hippocampus, bilateral pulvinar (lateral, medial and inferior parts), bilateral amygdala and left caudate whereas only the pulvinar was more activated by the Random condition (Supplementary Figure S1B right panel). Activations were stronger in the right superior colliculus (SC), right lateral geniculate nucleus (LGN), left caudate, left amygdala and in some portion of right and left pulvinar (lateral and inferior pulvinar) for ToM compared with Random animations (Supplementary Figure S1C right panel).

### Comparison of functional brain activations in humans and marmosets

As described above, compared to Random animations, ToM videos activated an extended network comprising a set of areas in occipital, temporal, parietal and frontal cortices in both humans and marmosets. Figure 5 shows human (A) and marmoset (B) flat maps of this comparison.

Overall, we observed a number of similarities between the two species, with comparable stronger activations for ToM animations in visual areas, in inferior and superior temporal areas, in the inferior parietal lobe and in several regions surrounding the IPS in the superior parietal lobe. We also found similar activations in somatosensory cortex, although more extended for marmosets than in humans, where activations were only located in secondary somatosensory cortex. Other similarities were found in premotor cortex and in some areas of the lateral prefrontal cortex. In general, left and right hemisphere activations were more similar in marmosets than in humans. However, this is likely due to our human head coil which had a lower SNR in the left hemisphere (see Supplementary Figure S3). Indeed, the human HCP dataset shows similar left and right activations in humans (Barch et al., 2013) (see Supplementary Figure S2B).

However, there were also some clear differences between the two species, including stronger activations in medial frontal cortex, primary motor area and posterior cingulate cortex for marmosets not observed in our human sample. Moreover, different parts of the insular cortex were recruited in marmosets, whereas in humans activations were limited to the parietal operculum and did not include the insula. Furthermore, at the subcortical level, more areas were activated in the ToM condition in marmosets, although amygdala and thalamic activations were present in humans and marmosets.

These results show that ToM animations were associated with stronger activations in both humans and marmosets. While there were many similarities in the activations, some clear differences were also observed.

## Discussion

In the present study, we investigated whether New-World common marmoset monkeys, like humans, process videos of animated abstract shapes differently when these move as if they are reacting to each other (ToM condition) compared to when they do not interact and move about purposelessly (Random condition). To directly compare the two primate species, we measured gaze patterns and brain activations while they viewed the popular Frith-Happé’s animations (Abell et al., 2000; Castelli et al., 2000). In the ToM animations, two triangles move as if one triangle is reacting to the other’s mental state (i.e., coaxing, surprising, seducing and mocking) whereas in the Random animations, the two triangles move independently in different patterns (i.e., billiard, drifting, star, tennis).

In our first experiment, we recorded the eye movements of marmosets and humans while subjects viewed the videos. Klein et al. (2009) reported longer fixation times for ToM compared to Random animations in human participants, interpreting them as the result of a mentalization effect. Indeed, eye movements measures and the use of fixation durations is known to provide a nonverbal measurement of the mentalizing capacity (Klein et al., 2009; Meijering et al., 2012). In contrast to Klein et al. (2009), however, Schafroth et al. (2021) did not find longer fixation duration for Frith-Happé’s ToM animations in macaque monkeys. Here, we confirmed the results of Klein et al. (2009) in humans, whereas marmosets, like macaques (Schafroth et al., 2021), did not show any significant difference between fixation durations in the two types of animations.

We then went further than the previous studies in humans (Klein et al., 2009) and macaques (Schafroth et al., 2021) and investigated the proportion of time subjects looked at each of the two triangles, the protagonists of the animations. While there was no difference in the proportion of time subjects looked at the large red and the small blue triangles during Random animations, both humans and marmosets spent significantly more time looking at the large red triangle during ToM compared with Random animations. In humans, but not marmosets, we also found the same, albeit weaker, effect for the small blue triangle.

Together, the eye tracking results do not provide support for the idea that marmosets increase their cognitive processing during ToM compared to random animations in the same fashion as humans. However, the findings clearly demonstrate that marmosets processed ToM animations differently than random animations as indicated by the increased time marmosets looked at the more salient large red triangle during the ToM animations.

Thus, in our second experiment we explored the brain networks associated with viewing the Frith-Happé’s animations in humans and marmosets. In humans, several fMRI studies have described a dedicated brain network for the processing of ToM stories or humorous cartoons involving complex mental states with activations mainly located in the medial frontal gyrus, the posterior cingulate, the inferior parietal cortex, and the temporoparietal junction (Fletcher et al., 1995; Gallagher et al., 2000). However, all these tasks involved complex stimuli and their findings were heterogeneous, implicating also various other brain regions (e.g. lateral prefrontal cortex, inferior parietal lobule, occipital cortex, insula) varying substantially between studies - as many experimental paradigms have been used to investigate ToM (Carrington and Bailey, 2009). A few studies have also used Frith-Happé’s animations, reporting a distinct pattern of brain activations in medial and lateral prefrontal cortex, inferior parietal cortex, temporoparietal, inferior and superior temporal regions as well as lateral superior occipital regions during the observation of ToM compared to Random animations. This is consistent with the idea that ToM animations - but not the Random ones – evoke mental state attributions (Barch et al., 2013; Castelli et al., 2000; Gobbini et al., 2007; Wheatley et al., 2007).

Here we confirmed that our slightly shortened versions of Frith-Happé’s animations elicit a similar distinct pattern of brain activations. Compared to HCP data in Barch et al. (2013), we found similar brain networks with preferential activations for ToM compared to Random animations in a set of areas in occipital, temporal, parietal and frontal cortices. Our results are similar to other previous fMRI studies (Barch et al., 2013; Castelli et al., 2000; Wheatley et al., 2007), although we did not find activations in the medial prefrontal cortex. In general, the ToM network in our study and in Barch et al. (2013) is more extended than what observed in the older studies (Castelli et al., 2000; Wheatley et al., 2007). Consistent with Barch et al. (2013), we found similar activations in visual areas, inferior and superior temporal areas including the STS, temporal parietal junction, posterior parietal cortex, lateral prefrontal cortex, and premotor cortex. At subcortical level, we found similar activations in cerebellum, thalamus and amygdala. The main difference in our study concerns wider activations in parietal cortex, involving the superior parietal lobule but also extended into somatosensory cortex. We also found fewer areas activated in the left compared to the right hemisphere, which is likely due to the lower SNR in our human coil on this side.

The comparison between ToM and Random animations in marmosets evoked responses in occipito-temporal, parietal and frontal areas. As in humans, these activations are located in dorsolateral prefrontal, premotor, secondary somatosensory, posterior parietal and visual cortices. In particular the activations in TE areas in marmosets could correspond to some of activations obtained along the STS in humans (Yovel and Freiwald, 2013). We also found some similar subcortical activations with our human subjects and previous human studies in amygdala, thalamus and caudate. As observed in the human literature (Carrington and Bailey, 2009; Fletcher et al., 1995; Gallagher et al., 2000), we also saw activations in inferior parietal cortex in both humans and marmosets. However, some differences between the results of our study and previous work are to be noted. First of all, both humans and marmosets involved in our study reported activations in the superior parietal cortex - in the area surrounding the IPS - which were not previously described (Barch et al., 2013; Carrington and Bailey, 2009; Fletcher et al., 1995; Gallagher et al., 2000). Furthermore, although we did not observe any activations in medial prefrontal cortex in humans, they were present in marmosets, a finding consistent with previous human fMRI studies (Castelli et al., 2000; Wheatley et al., 2007). The network in marmosets also included the posterior cingulate cortex and the insula, areas known to be involved respectively in mentalizing and affective processing in human ToM studies that used more complex stimuli (Fletcher et al., 1995; Gallagher et al., 2000; Wheatley et al., 2007). Finally, a prominent difference between humans and marmosets are the strong activations in marmoset motor cortex for ToM animations that are absent in humans. Interestingly, we recently also found activations in marmoset primary motor cortex during the observation of social interactions (Cléry et al., 2021), indicating a potential role for marmoset motor cortex in interaction observation. Together, these results demonstrate that the observation of interacting animated shapes compared to randomly moving shapes in marmosets is associated with stronger activation in a number of brain areas that have been previously associated with ToM processing in human subjects.

In summary, while our study cannot address the question of whether marmosets possess mentalization abilities similar to the humans’ theory of mind, the eye tracking results and fMRI activations both indicate that these New-World primates process moving abstract shapes differently when they are perceived to interact compared to when they move randomly. This clear preference for interacting shapes that we observe in the marmosets’ gaze patterns and in their cortical and subcortical activations may be an ancestral form or a prerequisite for the development of a theory of mind.

## Material and methods

### Common marmosets

All experimental procedures were in accordance with the Canadian Council of Animal Care policy and a protocol approved by the Animal Care Committee of the University of Western Ontario Council on Animal Care #2021-111.

Twelve adult marmosets (7 females, 32-57 months, mean age: 36.6 months) were subjects in this study. All animals were implanted for head-fixed experiments with either a fixation chamber (Johnston et al., 2018) or a head post (Gilbert et al., 2023) under anesthesia and aseptic conditions. Briefly, the animals were placed in a stereotactic frame (Narishige, model SR-6C-HT) while being maintained under gas anaesthesia with a mixture of O_2_ and air (isoflurane 0.5-3%). After a midline incision of the skin along the skull, the skull surface was prepared by applying two coats of an adhesive resin (All-Bond Universal; Bisco, Schaumburg, IL) using a microbrush, air-dried, and cured with an ultraviolet dental curing light (King Dental). Then, the head post or fixation chamber was positioned on the skull and maintained in place using a resin composite (Core-Flo DC Lite; Bisco). Heart rate, oxygen saturation, and body temperature were continuously monitored during this procedure.

Six of these animals (four females - weight 315-442 g, age 30-34 months - and two males - weight 374-425 g, age 30 and 55 months) were implanted with an MRI-compatible machined PEEK head post (Gilbert et al., 2023). Two weeks after the surgery, these marmosets were acclimatized to the head-fixation system in a mock MRI environment.

### Human participants

Twelve healthy humans (7 females, 23-54 years, mean age: 32.3 years) including three of the authors were subjects in the eye tracking experiment. Six of these subjects and four additional subjects (3 females, 25-54 years) participated in the fMRI experiment. All subjects self-reported as right-handed, had normal or corrected-to-normal vision and had no history of neurological or psychiatric disorders. Subjects were informed about the experimental procedures and provided informed written consent. These studies were approved by the Ethics Committee of the University of Western Ontario.

### Stimuli

Eight animations of simple geometric shapes with different movement patterns were used (Figure 1). These animations, originally developed by (Abell et al., 2000), showed two animated triangles, a big red triangle and a small blue one, moving in a way which indicates that one triangle reacts to the other object’s mental state (called ToM animations) or showing the same two triangles moving and bouncing like inanimate objects (called Random animations), on a framed white background. In more detail, in the ToM animations one triangle could (1) try to seduce and persuade the other to let it free, (2) mock the other one behind its back, (3) surprise the other one hiding behind a door, or (4) coax the other one out of an enclosure. In the Random animations, the two triangles didn’t interact with each other and moved independently in different patterns (billiard, drifting, star, tennis). As in the HCP (Barch et al., 2013), we used modified versions of these video clips. Each animation was shortened to 19.5 sec, instead of 40 sec, by truncating them using custom video-editing software (iMovie, Apple Incorporated, CA).

### Eye tracking task and data acquisition

To examine any behavioural differences while viewing TOM and Random animations, we presented all ToM and Random video clips once each in a pseudorandomized manner to both marmoset and human subjects. Stimulus presentation was controlled using Monkeylogic (Hwang et al., 2019). All stimuli were presented on a CRT monitor (ViewSonic Optiquest Q115, 76 Hz non-interlaced, 1600 × 1280 resolution). Eye position was digitally recorded at 1 kHz via video tracking of the left pupil (EyeLink 1000, SR Research, Ottawa, ON, Canada).

At the beginning of each session, horizontal and vertical eye positions of the left eye were calibrated by presenting a 1-degree dot at the display centre and at 6 degrees in each of the cardinal directions for 300 to 600ms. Monkeys were rewarded with a drop of diluted gum (50/50 mix of 1:1 acacia gum powder and water with liquid marshmallow) delivered via an infusion pump (model NE-510; New Era Pump Systems) through a liquid spout for successful fixations.

### fMRI experimental setup

During the scanning sessions, the marmosets sat in a sphinx position in a custom-designed plastic chair positioned within a horizontal magnet (see below). Their head was restrained using a head fixation system allowing to secure the surgically implanted head post to a clamping bar (Gilbert et al., 2023). After the head was immobilized, the two halves of the coil housing were positioned on either side of the head. Inside the scanner, monkeys faced a translucent screen placed 119 cm from their eyes where visual stimuli were projected with an LCSD-projector (Model VLP-FE40, Sony Corporation, Tokyo, Japan) via a back-reflection on a first surface mirror. Visual stimuli were presented with the Keynote software (version 12.0, Apple Incorporated, CA) and were synchronized with MRI TTL pulses triggered by a Raspberry Pi (model 3B+, Raspberry Pi Foundation, Cambridge, UK) running via a custom-written Python program. No reward was provided to the monkeys during the scanning sessions. Animals were monitored using an MRI-compatible camera (Model 12M-I, MRC Systems GmbH). Horizontal and vertical eye movements were monitored at 60Hz using a video eye tracker (ISCAN, Boston, Massachusetts). While we were able to obtain relatively stable eye movement recordings from a few runs per animal (min 1, max 5 runs per animal), the quality of the recordings was not sufficient for a thorough analysis. The large marmoset pupil represents a challenge for video eye tracking when the eyes are not fully open. Data from functional runs with more stable eye signals (n=15) show good compliance in the marmosets. The percentage of time spent in each run looking at the screen in the two experimental conditions (ToM, Random) and during the Baseline periods (fixation point in the center of the screen) was higher than 85% (88.2%, 88.6% and 93.4% respectively for ToM, Random and Baseline conditions). There was no significant differences between the ToM and Random condition (paired t-test, t_(14)_=-0.374, p=0.71), ruling out the possibility that any differences in fMRI activation between the ToM and Random condition were simply due to a different exposure to the videos.

Human subjects lay in a supine position and watched the stimuli presented via a rear projection system (Avotech SV-6011, Avotec Incorporated) through a surface mirror affixed to head coil. As for marmosets, visual stimuli were presented with the Keynote software (version 12.0, Apple Incorporated, CA) and were synchronized with MRI TTL pulses triggered by a Raspberry Pi (model 3B+, Raspberry Pi Foundation, Cambridge, UK) running via a custom-written python program.

### fMRI task

Humans and marmosets were presented with ToM and Random video clips in a block design. Each run consisted of eight blocks of stimuli (19.5 sec each) interleaved by baseline blocks (15 sec each). ToM or Random animations were presented pseudorandomly, and each condition was repeated four times (Figure 1). For each run, the order of these conditions was randomized leading to 14 different stimulus sets, counterbalanced within and between subjects. In baseline blocks, a 0.36° circular black cue was displayed at the center of the screen against a gray background. We found previously that such a stimulus reduced the vestibulo-ocular reflex evoked by the strong magnetic field.

### MRI data acquisition

Marmoset and human imaging were performed at the Center for Functional and Metabolic Mapping at the University of Western Ontario.

For marmoset subjects, fMRI data were acquired on a 9.4T 31 cm horizontal bore magnet (Varian) with a Bruker BioSpec Avance III HD console running software package Paravision-360 (Bruker BioSpin Corp), a custom-built high-performance 15-cm diameter gradient coil (maximum gradient strength: 1.5 mT/m/A), and an eight-channel receive coil. Preamplifiers were located behind the animals, and the receive coil was placed inside an in-house built quadrature birdcage coil (12-cm inner diameter) used for transmission. Functional images were acquired during 6 functional runs for each animal using gradient-echo based single-shot echo-planar images (EPI) sequence with the following parameters: TR=1.5s, TE = 15ms, flip angle = 40°, field of view=64×48 mm, matrix size = 96×128, resolution of 0.5 mm3 isotropic, number of slices= 42 [axial], bandwidth=400 kHz, GRAPPA acceleration factor: 2 (left-right). Another set of EPIs with an opposite phase-encoding direction (right-left) was collected for the EPI-distortion correction. A T2-weighted structural was also acquired for each animal during one of the sessions with the following parameters: TR=7s, TE=52ms, field of view=51.2×51.2 mm, resolution of 0.133×0.133×0.5 mm, number of slices= 45 [axial], bandwidth=50 kHz, GRAPPA acceleration factor: 2.

For human subjects, fMRI data were acquired on a 7T 68 cm MRI scanner (Siemens Magnetom 7T MRI Plus) with an AC-84 Mark II gradient coil, an in-house 8-channel parallel transmit, and a 32-channel receive coil (Gilbert et al., 2021). Functional images were acquired during 3 functional runs for each participant using Multi-Band EPI BOLD sequences with the following parameters: TR=1.5s, TE = 20ms, flip angle = 30°, field of view=208×208 mm, matrix size = 104×104, resolution of 2 mm3 isotropic, number of slices= 62, GRAPPA acceleration factor: 3 (anterior-posterior), multi-band acceleration factor: 2. Field map images were also computed from the magnitude image and the two phase images. An MP2RAGE structural image was also acquired for each subject during the sessions with the following parameters: TR=6s, TE=2.13 ms, TI1 / TI2 = 800 / 2700 ms, field of view=240×240 mm, matrix size= 320×320, resolution of 0.75 mm3 isotropic, number of slices= 45, GRAPPA acceleration factor (anterior posterior): 3.

### MRI data preprocessing

Marmoset fMRI data were preprocessed using AFNI (Cox, 1996) and FSL (Smith et al., 2004) software packages. Raw MRI images were first converted to NIfTI format using dcm2nixx AFNI’s function and then reoriented to the sphinx position using fslswapdim and fslorient FSL’s functions. Functional images were despiked using 3Ddespike AFNI’s function and time shifted using 3dTshift AFNI’s function. Then, the images obtained were registered to the base volume (i.e., corresponding to the middle volume of each time series) with 3dvolreg AFNI’s function. The output motion parameters obtained from volume registration were later used as nuisance regressors. All fMRI images were spatially smoothed with a 1.5 mm half-maximum Gaussian kernel (FWHM) with 3dmerge AFNI’s function, followed by temporal filtering (0.01-0.1 Hz) using 3dBandpass AFNI’s function. The mean functional image was calculated for each run and linearly registered to the respective anatomical image of each animal using FMRIB’s linear registration tool (FLIRT).

The transformation matrix obtained after the registration was then used to transform the 4D time series data. The brain was manually skull-stripped from individual anatomical images using FSL eyes tool and the mask of each animal was applied to the functional images. Finally, the individual anatomical images were linearly registered to the NIH marmoset brain template (Liu et al., 2018) using Advanced Normalization Tools (ANTs).

Human fMRI data were preprocessed using SPM12 (Wellcome Department of Cognitive Neurology). After converting raw images into NifTI format, functional images were realigned to correct for head movements and underwent slice timing correction. A field map correction was applied to the functional images from the magnitude and phase images with the specify toolbox implemented in SPM. Then, the anatomical and functional volumes corrected were coregistered with the MP2RAGE structural scan from each individual participant and normalized to the Montreal Neurological Institute (MNI) standard brain space. Anatomical images were segmented into white matter, gray matter, and CSF partitions and also normalized to the MNI space. The functional images were then spatially smoothed with a 6 mm FWHM isotropic Gaussian kernel. A high-pass filter (128 s) was also applied to the time series.

### Statistical analysis

#### Behavioral eye tracking data

To evaluate gaze patterns during observation of ToM and Random videos, we used mixed analyses of variance (ANOVA), with factors of species (Human vs Marmoset) and condition (ToM vs Random videos) on fixation duration in general and on the proportion of time when the radial distance between the subject’s gaze position and each triangle was less than 4 degrees. Partial eta squared (*η*_*p*_^*2*^) was computed as a measure of effect size and *post-hoc* comparisons were Bonferroni corrected.

#### fMRI data

For each run, a general linear regression model was defined: the task timing was convolved to the hemodynamic response (AFNI’s ‘BLOCK’ convolution for marmosets’ data and SPM12 hemodynamic response function for humans’ data) and a regressor was generated for each condition (AFNI’s 3dDeconvolve function for marmosets and SPM12 function for humans). The two conditions were entered into the same model, corresponding to the 19.5 sec presentation of the stimuli, along with polynomial detrending regressors and the marmosets’ motions parameters or human’s head movement parameters estimated during realignment.

The resultant regression coefficient maps of marmosets were then registered to template space using the transformation matrices obtained with the registration of anatomical images on the template (see MRI data processing part above).

Finally, we obtained for each run in marmosets and humans, two T-value maps registered to the NIH marmoset brain atlas (Liu et al., 2018) and to the MNI brain standard space, respectively.

These maps were then compared at the group level via paired t-tests using AFNI’s 3dttest++ function, resulting in Z-value maps. To protect against false positives and to control for multiple comparisons, we adopted a clustering method derived from 10000 Monte Carlo simulations to the resultant z-test maps using ClustSim option (α=0.05). This method corresponds to performing cluster-forming threshold of p<0.01 uncorrected and then applying a family-wise error (FWE) correction of p<0.05 at the cluster-level.

We used the Paxinos parcellation of the NIH marmoset brain atlas (Liu et al., 2018) and the most recent multi-modal cortical parcellation atlas (Glasser et al., 2016) to define anatomical locations of cortical and subcortical regions for both marmosets and humans respectively.

We first identified brain regions involved in the processing of ToM and Random animations separately (i.e., ToM condition > baseline and Random condition > baseline contrasts). We then examined the clusters that were significantly more activated by ToM compared to Random animations (ToM condition > Random condition contrast), and vice versa. The resultant Z-value maps were displayed on fiducial maps obtained from the Connectome Workbench (v1.5.0 (Marcus et al., 2011)) using the NIH marmoset brain template (Liu et al., 2018) for marmosets and the MNI Glasser brain template (Glasser et al., 2016) for humans. Subcortical activations were displayed on coronal sections.

As we used shortened video clips (i.e. 19.5 sec compared to the 40 sec originally designed by Abell et al., 2000), we validated our fMRI protocol by confirming that our shorter videos elicited similar responses to those previously observed in the HCP (Barch et al., 2013), which is the only study that also used modified versions of these animation videos. We compared our ToM vs Random Z-value map obtained in human subjects with those of the HCP (Barch et al., 2013). To this end, we downloaded the Z-value map of activations for ToM animations compared to Random animations from 496 subjects from the Neurovalt site (https://identifiers.org/neurovault.image:3179). We displayed the resultant Z-value maps on fiducial maps obtained from the Connectome Workbench (v1.5.0, (Marcus et al., 2011)) using the MNI Glasser brain template (Glasser et al., 2016).

## Acknowledgements

Support was provided by the Canadian Institutes of Health Research (FRN 148365), the Canada First Research Excellence Fund to BrainsCAN, and a Discovery grant by the Natural Sciences and Engineering Research Council of Canada. We are grateful to Drs. Sarah White and Ulla Frith for access to the Frith-Happé animation videos. We also wish to thank Cheryl Vander Tuin, Whitney Froese, Hannah Pettypiece, and Miranda Bellyou for animal preparation and care, Dr. Alex Li and Trevor Szekeres for scanning assistance, Dr. Kyle Gilbert and Peter Zeman for coil designs.

## Author contributions

A.D., A.Z., and S.E. designed research. A.D., A.Z., and J.S. performed research and analysed data. A.D. wrote the manuscript. A.D., A.Z, J.S., R.M., and S.E. edited the manuscript.

## References

Abell F, Happé F, Frith U. 2000. Do triangles play tricks? Attribution of mental states to animated shapes in normal and abnormal development. Cogn Dev 15:1–16. doi:10.1016/S0885-2014(00)00014-9.

Atsumi T, Koda H, Masataka N. 2017. Goal attribution to inanimate moving objects by Japanese macaques (Macaca fuscata). Sci Reports 2017 71 7:1–7. doi:10.1038/srep40033.

Atsumi T, Nagasaka Y. 2015. Perception of chasing in squirrel monkeys (Saimiri sciureus). Anim Cogn 18:1243–1253. doi:10.1007/S10071-015-0893-X/FIGURES/5.

Barch DM, Burgess GC, Harms MP, Petersen SE, Schlaggar BL, Corbetta M, Glasser MF, Curtiss S, Dixit S, Feldt C, Nolan D, Bryant E, Hartley T, Footer O, Bjork JM, Poldrack R, Smith S, Johansen-Berg H, Snyder AZ, Van Essen DC. 2013. Function in the Human Connectome: Task-fMRI and Individual Differences in Behavior. Neuroimage 80:169. doi:10.1016/J.NEUROIMAGE.2013.05.033.

Bowler DM, Thommen E. 2000. Attribution of mechanical and social causality to animated displays by children with autism. SAGE Publ Natl Autistic Soc 4:1362–3613.

Burkart JM, Finkenwirth C. 2015. Marmosets as model species in neuroscience and evolutionary anthropology. Neurosci Res 93:8–19. doi:10.1016/J.NEURES.2014.09.003.

Burkart JM, Hrdy SB, Van Schaik CP. 2009. Cooperative breeding and human cognitive evolution. Evol Anthropol Issues, News, Rev 18:175–186. doi:10.1002/EVAN.20222.

Carrington SJ, Bailey AJ. 2009. Are there theory of mind regions in the brain? A review of the neuroimaging literature. Hum Brain Mapp 30:2313. doi:10.1002/HBM.20671.

Carruthers P, Smith PK. 1996. Theories of Theories of Mind.

Castelli F, Frith C, Happé F, Frith U. 2002. Autism, Asperger syndrome and brain mechanisms for the attribution of mental states to animated shapes. Brain 125:1839–1849. doi:10.1093/BRAIN/AWF189.

Castelli F, Happé F, Frith U, Frith C. 2000. Movement and mind: a functional imaging study of perception and interpretation of complex intentional movement patterns. Neuroimage 12:314–325. doi:10.1006/NIMG.2000.0612.

Cléry JC, Hori Y, Schaeffer DJ, Menon RS, Everling S. 2021. Neural network of social interaction observation in marmosets. Elife 10. doi:10.7554/ELIFE.65012.

Cox RW. 1996. AFNI: Software for analysis and visualization of functional magnetic resonance neuroimages. Comput Biomed Res 29:162–173. doi:10.1006/cbmr.1996.0014.

Fletcher PC, Happé F, Frith U, Baker SC, Dolan RJ, Frackowiak RSJ, Frith CD. 1995. Other minds in the brain: a functional imaging study of “theory of mind” in story comprehension. Cognition 57:109–128. doi:10.1016/0010-0277(95)00692-R.

Gallagher HL, Happé F, Brunswick N, Fletcher PC, Frith U, Frith CD. 2000. Reading the mind in cartoons and stories: an fMRI study of ‘theory of mind’ in verbal and nonverbal tasks. Neuropsychologia 38:11–21. doi:10.1016/S0028-3932(99)00053-6.

Gilbert KM, Dureux A, Jafari A, Zanini A, Zeman P, Menon RS, Everling S. 2023. A radiofrequency coil to facilitate task-based fMRI of awake marmosets. J Neurosci Methods 383:109737. doi:10.1016/J.JNEUMETH.2022.109737.

Gilbert KM, Klassen LM, Mashkovtsev A, Zeman P, Menon RS, Gati JS. 2021. Radiofrequency coil for routine ultra-high-field imaging with an unobstructed visual field. NMR Biomed 34. doi:10.1002/NBM.4457.

Glasser MF, Smith SM, Marcus DS, Andersson J, Auerbach EJ, Behrens TEJ, Coalson TS, Harms MP, Jenkinson M, Moeller S, Robinson EC, Sotiropoulos SN, Xu J, Yacoub E, Ugurbil K, Essen DC Van. 2016. The Human Connectome Project’s Neuroimaging Approach. Nat Neurosci 19:1175. doi:10.1038/NN.4361.

Gobbini MI, Koralek AC, Bryan RE, Montgomery KJ, Haxby J V. 2007. Two Takes on the Social Brain: A Comparison of Theory of Mind Tasks. J Cogn Neurosci 19:1803–1814. doi:10.1162/JOCN.2007.19.11.1803.

Happé FGE. 1994. An advanced test of theory of mind: understanding of story characters’ thoughts and feelings by able autistic, mentally handicapped, and normal children and adults. J Autism Dev Disord 24:129–154. doi:10.1007/BF02172093.

Heider F, Simmel M. 1944. An Experimental Study of Apparent Behavior. Am J Psychol 57:243. doi:10.2307/1416950.

Hwang J, Mitz AR, Murray EA. 2019. NIMH MonkeyLogic: Behavioral control and data acquisition inMATLAB. J Neurosci Methods 323:13. doi:10.1016/J.JNEUMETH.2019.05.002.

Johnston KD, Barker K, Schaeffer L, Schaeffer D, Everling S. 2018. Methods for chair restraint and training of the common marmoset on oculomotor tasks. J Neurophysiol 119:1636–1646. doi:10.1152/JN.00866.2017/SUPPL_FILE/SUPPLEMENTAL.

Klein AM, Zwickel J, Prinz W, Frith U. 2009. Animated triangles: An eye tracking investigation. Q J Exp Psychol 62:1189–1197. doi:10.1080/17470210802384214/ASSET/IMAGES/LARGE/10.1080_17470210802384214-FIG2.JPEG.

Klin A. 2000. Attributing Social Meaning to Ambiguous Visual Stimuli in Higher-functioning Autism and Asperger Syndrome: The Social Attribution Task. J Child Psychol Psychiatry 41:831–846. doi:10.1111/1469-7610.00671.

Krupenye C, Hare B. 2018. Bonobos Prefer Individuals that Hinder Others over Those that Help. Curr Biol 28:280-286.e5. doi:10.1016/J.CUB.2017.11.061.

Kupferberg A, Glasauer S, Burkart JM. 2013. Do robots have goals? How agent cues influence action understanding in non-human primates. Behav Brain Res 246:47–54. doi:10.1016/J.BBR.2013.01.047.

Liu C, Ye FQ, Yen CCC, Newman JD, Glen D, Leopold DA, Silva AC. 2018. A digital 3D atlas of the marmoset brain based on multi-modal MRI. Neuroimage 169:106–116. doi:10.1016/J.NEUROIMAGE.2017.12.004.

Marcus DS, Harwell J, Olsen T, Hodge M, Glasser MF, Prior F, Jenkinson M, Laumann T, Curtiss SW, Van Essen DC. 2011. Informatics and data mining tools and strategies for the human connectome project. Front Neuroinform 5:4. doi:10.3389/FNINF.2011.00004/BIBTEX.

Meijering B, van Rijn H, Taatgen NA, Verbrugge R. 2012. What Eye Movements Can Tell about Theory of Mind in a Strategic Game. PLoS One 7:e45961. doi:10.1371/JOURNAL.PONE.0045961.

Miller CT, Freiwald WA, Leopold DA, Mitchell JF, Silva AC, Wang X. 2016. Marmosets: A Neuroscientific Model of Human Social Behavior. Neuron 90:219. doi:10.1016/J.NEURON.2016.03.018.

Premack D, Woodruff G. 1978. Does the chimpanzee have a theory of mind? Behav Brain Sci 1:515–526. doi:10.1017/S0140525×00076512.

Sarfati Y, Hardy-Baylé MC, Besche C, Widlöcher D. 1997. Attribution of intentions to others in people with schizophrenia: a non-verbal exploration with comic strips. Schizophr Res 25:199–209. doi:10.1016/S0920-9964(97)00025-X.

Schafroth JL, Basile BM, Martin A, Murray EA. 2021. No evidence that monkeys attribute mental states to animated shapes in the Heider–Simmel videos. Sci Reports 2021 111 11:1–10. doi:10.1038/s41598-021-82702-6.

Smith SM, Jenkinson M, Woolrich MW, Beckmann CF, Behrens TEJ, Johansen-Berg H, Bannister PR, De Luca M, Drobnjak I, Flitney DE, Niazy RK, Saunders J, Vickers J, Zhang Y, De Stefano N, Brady JM, Matthews PM. 2004. Advances in functional and structural MR image analysis and implementation as FSL. Neuroimage 23:S208–S219. doi:10.1016/J.NEUROIMAGE.2004.07.051.

Uller C. 2004. Disposition to recognize goals in infant chimpanzees. Anim Cogn 7:154–161. doi:10.1007/S10071-003-0204-9/TABLES/2.

Wheatley T, Milleville SC, Martin A. 2007. Understanding animate agents: Distinct roles for the social network and mirror system: Research report. Psychol Sci 18:469–474. doi:10.1111/j.1467-9280.2007.01923.x.

Wimmer H, Perner J. 1983. Beliefs about beliefs: Representation and constraining function of wrong beliefs in young children’s understanding of deception. Cognition 13:103–128. doi:10.1016/0010-0277(83)90004-5.

Yovel G, Freiwald WA. 2013. Face recognition systems in monkey and human: are they the same thing? F1000Prime Rep 5. doi:10.12703/P5-10.

