## Supplementary figures for "Gaze patterns and brain activations in humans and marmosets in the Frith-Happé theory-of-mind animation task"

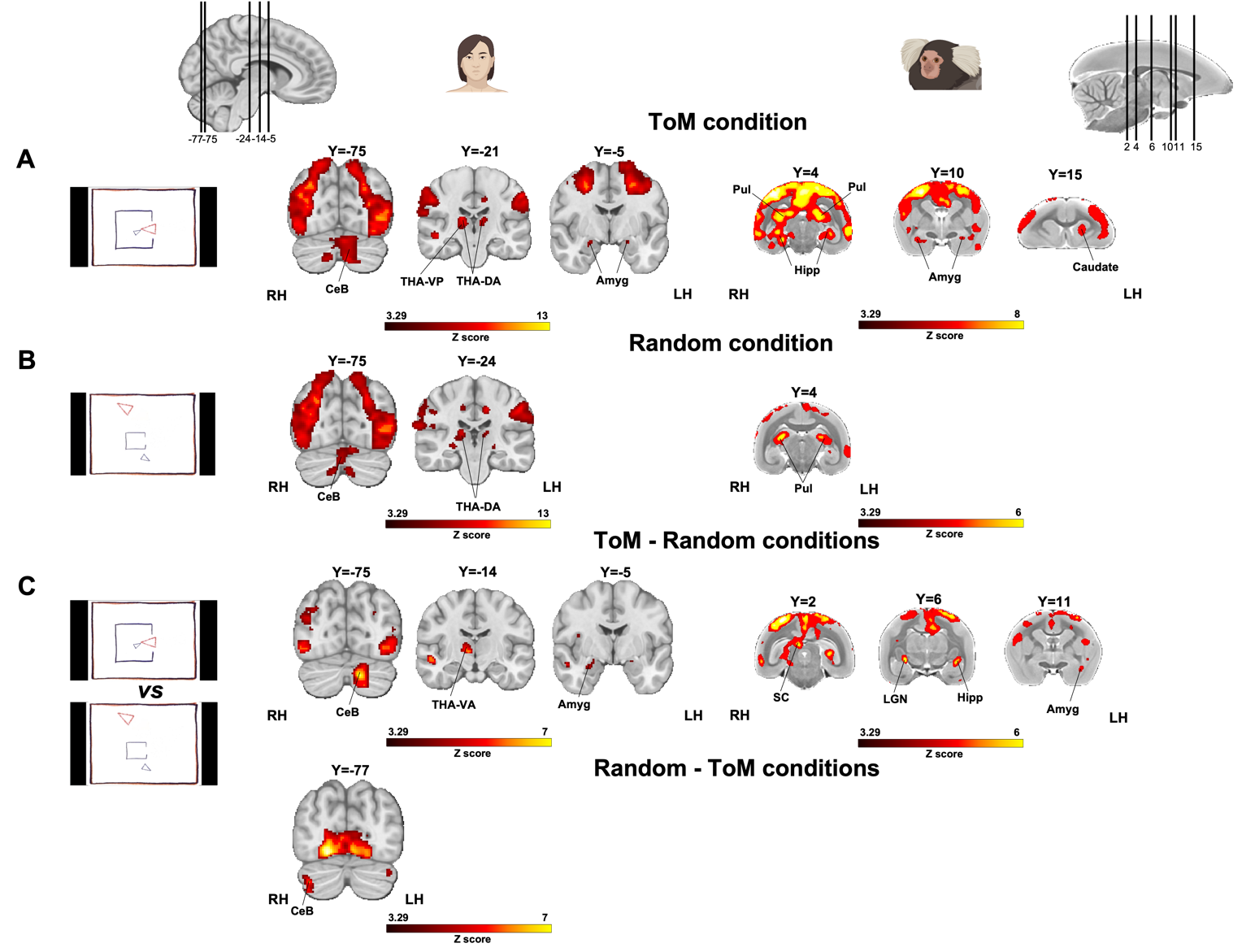

**Figure S1*.* Subcortical activations during processing of Frith-Happé’s ToM and Random animations in humans (left) and marmosets (right).** Group subcortical functional maps showing significant greater activations for ToM condition (A), Random condition (B) and the comparison between ToM and Random conditions (C). Group map displayed on coronal slices obtained from ten humans (left side) and 6 marmosets (right side). The brain areas reported have activation threshold corresponding to z-scores > 3.29 (AFNI’s 3dttest++, threshold of p<.001 uncorrected). *CeB, cerebellum; THA-VP, ventroposterior thalamus; THA-DA, dorsoanterior thalamus; THA-VA, ventroanterior thalamus; Amyg, amygdala; Hipp, hippocampus; Pul, pulvinar; SC, superior colliculus; LGN, lateral geniculate nucleus.*

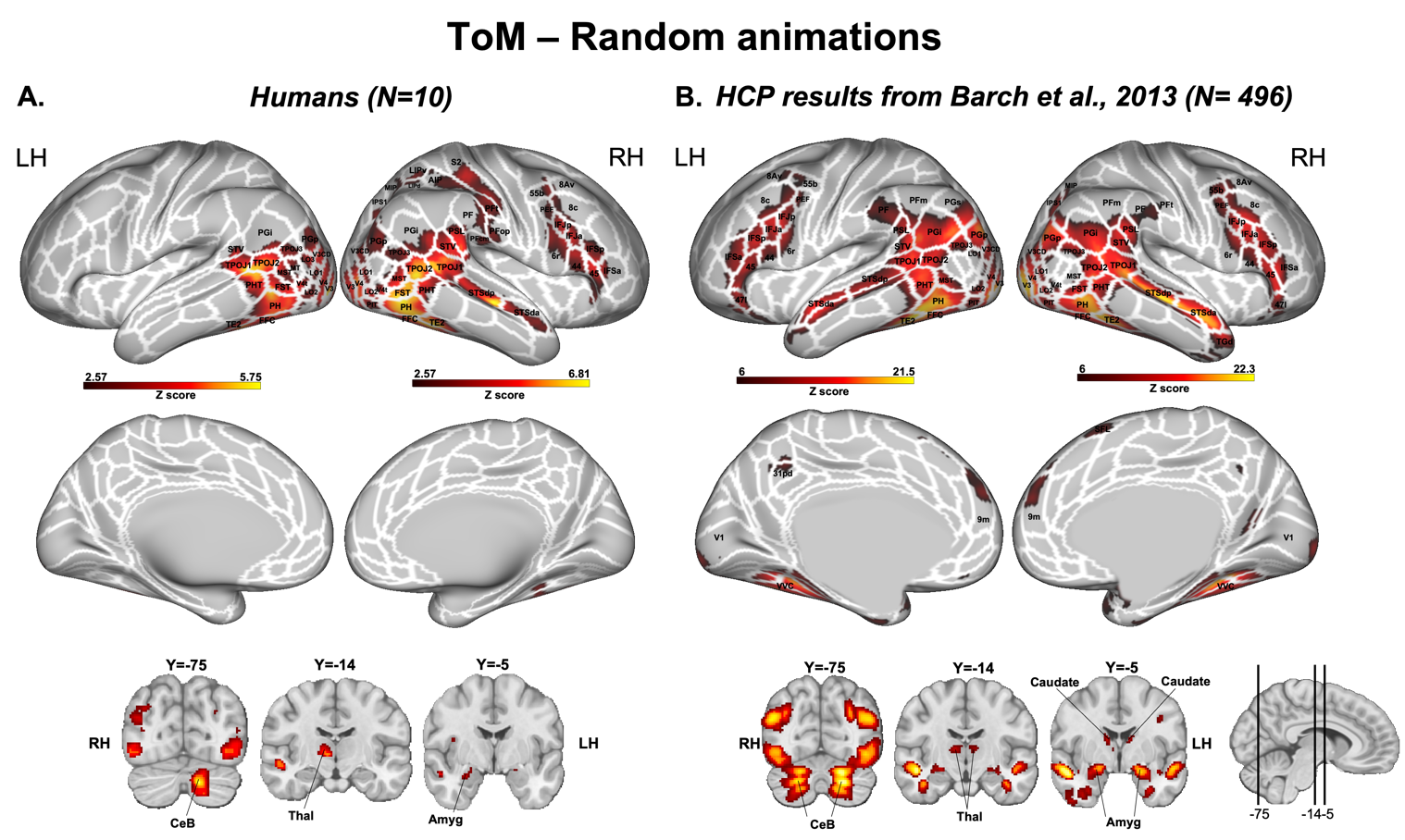

**Figure S2*.* Brain networks involved in ToM animations processing in humans.** Group functional maps showing significant greater activations for the comparison between ToM animations and Random animations displayed on the left and right fiducial human cortical surfaces (lateral and medial views) as well as on coronal slices, to illustrate the activations in subcortical areas. The white line delineates the regions based on the recent multi-modal cortical parcellation atlas ^34^. **A.** The map depicted is obtained from 10 human subjects with an activation threshold corresponding to z-scores *>* 2.57 (AFNI’s 3dttest++, cluster-forming threshold of p<.01 uncorrected and then FWE-corrected α=.05 at cluster-level from 10000 Monte-Carlo simulations). The subcortical maps correspond to an activation threshold of z-scores > 3.29 (AFNI’s 3dttest++, threshold of p<0.001 uncorrected). **B.** The map depicted has been downloaded from <https://identifiers.org/neurovault.image:3179> and is described in the study of Barch et al. (2013). The brain areas reported have activation threshold corresponding to z-scores > 6, uncorrected.

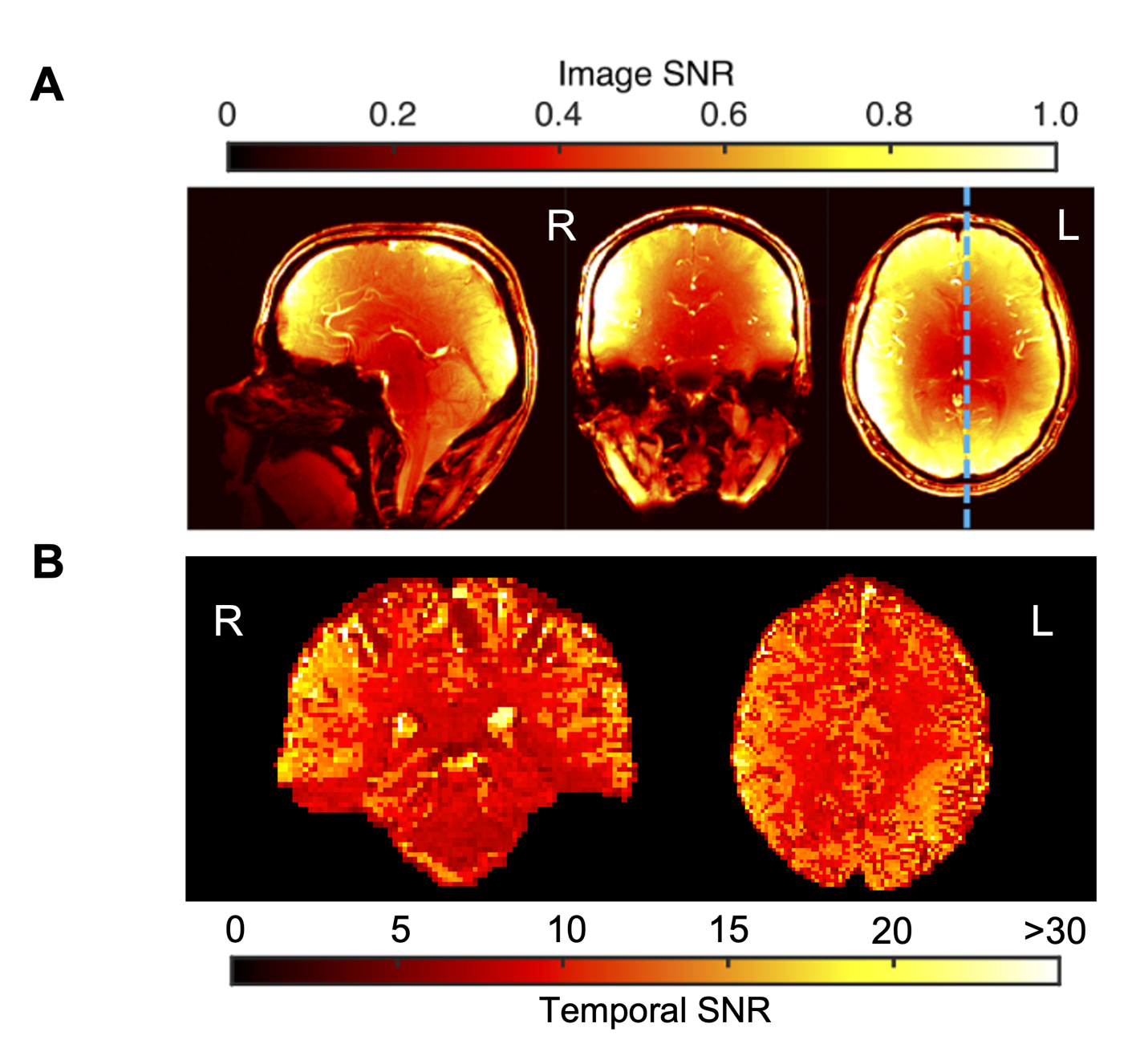

**Figure S3. Image and temporal signal-to-noise-ration (SNR) calculated on fMRI data acquired at 7T with an AC-84 Mark II gradient coil, an in-house 8-channel transmit, and a 32-channel receive coil (see methods, main text). A.** Image SNR maps from gradient-echo-images obtained from Gilbert et al., 2021. **B.** Temporal SNR (i.e., ratio of the mean signal to the standard deviation through the time course) for EPI BOLD images obtained from one run of one participant. The mean tSNR calculated within peripheral brain regions nearest the coil elements is 10% higher for right hemisphere than tSNR produced by left hemisphere.
